# Multiple transcription factors regulate the expression of genes for error prone DNA polymerases in *Acinetobacter baumannii*

**DOI:** 10.1101/2024.09.30.615892

**Authors:** Brian Nguyen, Carly Ching, Ashley Macguire, Pranav Casula, Connor Newman, Faith Finley, Veronica G Godoy

## Abstract

*Acinetobacter baumannii* is an opportunistic pathogen causing several infections that are increasingly difficult to treat due to its ability to rapidly gain antibiotic resistances. These resistances can arise due to mutations during the DNA damage response (DDR), through the activity of error-prone DNA polymerases, such as DNA polymerase V (DNA Pol V). Currently, the DDR in *A. baumannii* is not well understood and the regulation of genes encoding multiple copies of DNA Pol V is not fully characterized. Through genome wide mutagenesis, we have identified a novel TetR-like family regulator of genes that encode DNA Pol V, which we have named <u>E</u>rror-<u>p</u>rone <u>p</u>olymerase regulator, EppR. We have found that EppR represses the expression of the genes encoding DNA Pol V and itself through direct binding to a conserved EppR motif in their promoters. Lastly, we show that EppR also regulates UmuDAb, previously identified as a regulator of genes encoding DNA Pol V. These two gene products are functionally required to ensure regulation of expression of genes encoding DNA Pol Vs and *umuDAb*. With these results, we propose a co-repressor model for the regulation of genes encoding DNA Pol V and *umuDAb*.

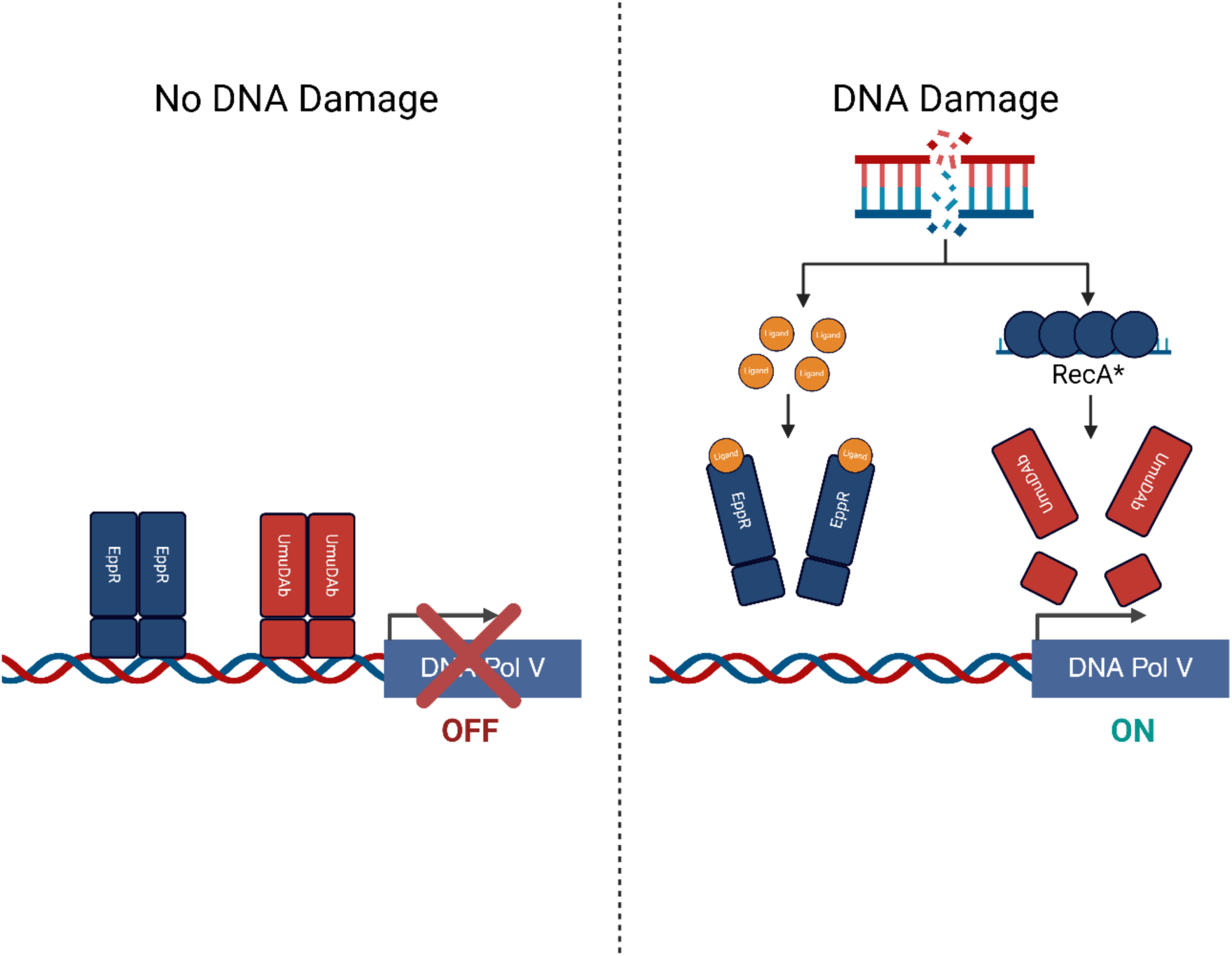
Graphical Abstract.

## Introduction

In many bacterial species, extensive DNA damage is resolved by the well-known SOS response or DNA damage response (DDR). The current understanding of bacterial DDRs is based of *Escherichia coli*, where DNA damage leads to the accumulation of single-stranded DNA (ssDNA) which is then bound by the recombinase RecA to form the RecA nucleoprotein filament (RecA*), which in turns interacts with the DDR master regulator, LexA, to facilitate its autocleavage. LexA is a repressor that binds to a common binding motif present in all DDR-regulated genes; its autocleavage results in DDR activation(Butala et al., n.d.; Janion, 2008; Little and Mount, 1982). This strategy for dealing with DNA damage has been found to be highly conserved in other bacterial species(Cirz et al., 2007; Ryan T Cirz et al., 2006; Sanchez-Alberola et al., 2012). In contrast to *E. coli*, *Acinetobacter baumannii* does not share a LexA homologue and genes that traditionally make up the DDR do not seem to share a common regulatory motif, suggesting that multiple different regulators may be involved in DDR regulation(Robinson et al., 2010). Because of these differences, *A. baumannii* is considered to have a noncanonical DDR. While *A. baumannii* does not have a LexA homologue, previous work in our lab has shown that DDR genes still respond to DNA damage in an organized fashion (Norton et al., 2013). In response to DNA damage, classical DNA response genes (e.g., *recA*, *uvrA*) display differential expression, where subpopulations of both high and low DDR-expressing cells are present (MacGuire et al., 2014). Additional work in the lab has provided insight into how *A. baumannii recA*, a key master DDR regulator, governs its regulation and the response to DNA damage (Ching et al., 2017). The regulation of other DDR genes, including error-prone DNA polymerases, and how they link to RecA continues to remain unclear. We had previously shown that in *A. baumannii* there is a causal relationship between the DDR and antibiotic-resistance acquisition, where we found an increase in antibiotic-resistant mutants upon DDR induction (Norton et al., 2013). This relationship has been known in other bacterial species and studies have shown that the activity of error-prone DNA polymerases, namely DNA polymerase V (DNA Pol V) has led to increases in antibiotic-resistance acquisition (Boshoff et al., 2003; Cirz et al., 2005; Ryan T. Cirz et al., 2006; Cirz and Romesberg, 2006). Due to ease of which *A. baumannii* can gain antibiotic resistances (Bergogne-Bérézin and Towner, 1996), it is crucial for us to understand DDR regulation, especially the regulation of DNA Pol V genes.

DNA Pol V is encoded by the genes *umuDC* and its function is to synthesize DNA over a wide variety of DNA lesions, though with low fidelity. This is one of the drivers of bacterial mutagenesis in response to DNA damage (Janion, 2008; Little and Mount, 1982; Wang, 2001). Interestingly, the *A. baumannii* strain 17978 contains multiple copies of the *umuDC* operon, *umuC*, and a single additional copy of *umuD* while *E. coli* only has a single *umuDC* operon (Norton et al., 2013). This suggests that *A. baumannii* has a higher capacity for DNA-damage induced mutagenesis of drug targets, that could contribute to antibiotic resistance. Other groups have shown that these *umuDC* genes are regulated by UmuD^1389^, also known as UmuDAb (Aranda et al., 2013; Hare et al., 2006a; Witkowski et al., 2016). Although UmuDAb is homologous to the UmuD subunit of Pol V in *E. coli*, UmuDAb contains an additional DNA-binding domain in its N-terminal domain. This has not been found in other UmuD proteins, even in *A. baumannii* (Aranda et al., 2014; Hare et al., 2006b). With this DNA-binding domain, UmuDAb acts as a transcriptional repressor that recognizes and binds to a conserved binding motif present in *umuDC* promoters (Aranda et al., 2014). It has also been found that truncated forms of UmuDAb do not repress *umuDC* expression *in vivo* (Witkowski et al., 2016), and that UmuDAb is cleaved in response to DNA damage (Hare et al., 2012), providing a regulatory mechanism. Previous work in our lab had also identified the UmuDAb binding motif. We found that scrambling on the chromosome the UmuDAb binding motif sequence in one of the *umuDC* promoters resulted in either only high or low expressing cells of a chromosomal fluorescent *umuDC* reporter depending on the DNA damaging treatment compared to the parental (MacGuire et al., 2014). There are also conflicting data on whether UmuDAb is a repressor or an activator of certain DNA Pol V encoding genes (Aranda et al., 2013; Hare et al., 2014).These data brought up the possibility that there may be additional regulators to explain the data.

To test the hypothesis, we designed an experiment to isolate additional DNA Pol V gene regulators after random mutagenesis of the GFP-*umuDC* reporter strain (MacGuire et al., 2014). Using this approach, we identified a novel TetR-like transcriptional regulator of DNA Pol V genes. We present evidence to support a repressor function for this *A. baumannii* gene product through a combination of *in vivo* and *in vitro* assays. In addition, we present evidence showing that this novel regulator acts with UmuDAb to coregulate DNA Pol V expression.

## Results

### Methyl Methanesulfonate (MMS) Mutagenesis and Identification of EppR

To identify potential regulators of DNA polymerase V (DNA Pol V) genes, we first constructed a chromosomal GFP transcriptional reporter with one of the *umuDC* promoters (*umuDC^0636-0637^*) (MacGuire et al., 2014) and used a genome wide approach by treating cells with methyl methanesulfonate (MMS), an alkylating agent, as we have previously done (Norton et al., 2013). After treatment, we grew cells without MMS on LB Agar plates and identified colonies that showed a difference in fluorescence intensity with and without DNA damage treatment (Figure 1). We expected that cells with mutations on a potential regulator would show differences in fluorescence intensity compared to MMS-untreated cells.

**Figure 1:**
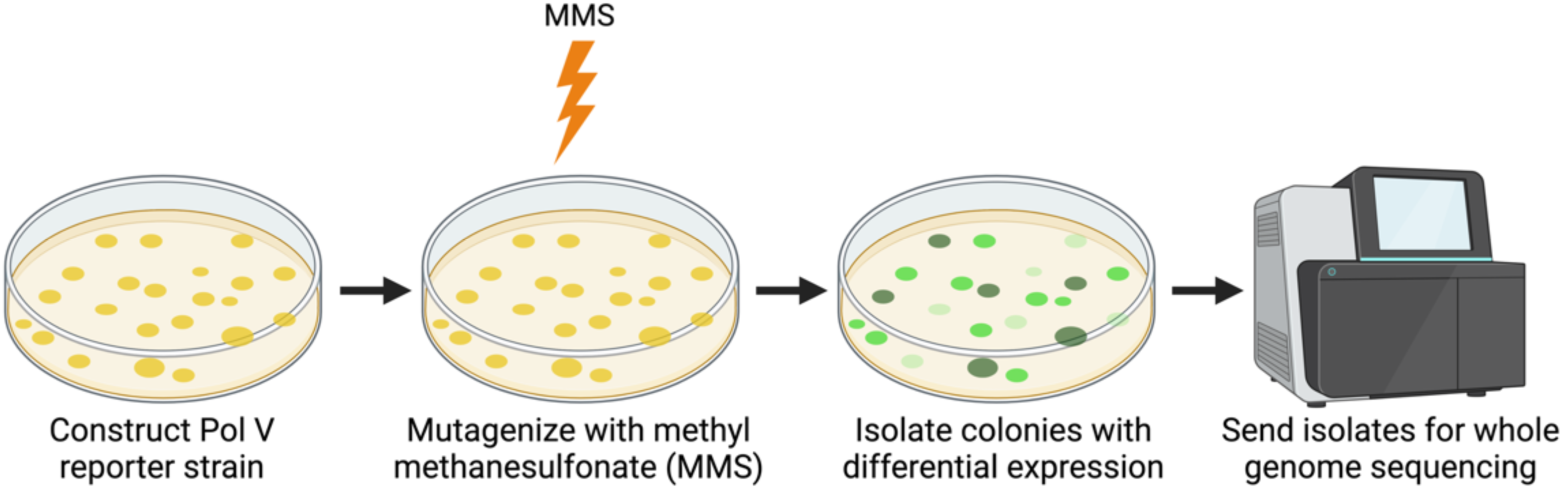
Seeking a DDR regulator by random mutagenesis with methyl methanesulfonate (MMS). To identify novel regulators of genes encoding DNA Pol V, a chromosomal *umuDC^0636-0637^* transcriptional GFP reporter strain was constructed (A. baumannii *umuDC-GFP*, Table S1) and treated with a sublethal concentration of methyl methanesulfonate (MMS). Colonies that exhibit high GFP signal (represented by the dark green circles in the diagram) in the absence of DNA damage indicate a possible mutation in a potential repressor of genes encoding DNA Pol V. Low GFP signal during DNA damage (represented by the light green circles on the plates in the diagram) indicate a possible mutation in a potential activator. Colonies that exhibit differential expression were isolated and sent for whole genome sequencing. Figure created with biorender.com.

Out of approximately 10,000 colonies screened with a low magnification fluorescent microscopy, we identified 18 colonies that were highly fluorescent in the absence of DNA damage treatment. To ensure that the increased expression wasn’t due to mutations either in *gfp* or its promoter, we amplified and sequenced the chromosomal reporter. We found that all bright mutants did not have mutations within the reporter fragment. We also sequenced the *umuD^1389^* gene (A1S_1389; ACX60_RS11175) as it encodes UmuDAb, which is known to regulate genes that code for DNA Pol V in *A. baumannii* (Aranda et al., 2013; Hare et al., 2014). None of the mutant strains contained a mutation within this gene. Afterwards, candidates were sent for Illumina whole genome sequencing.

Of the mutant strains tested, we focused on mutant 1, which we will refer to as Bright Mutant 1 (BM1, Table S1), as it was the only isolate that contained a mutation resulting in a stop codon. This nonsense mutation was located within the open reading frame (ORF) of a putative TetR-like family regulator (A1S_1669, ACX60_RS09590). Moreover, this strain had two missense mutations within predicted ORFs, one in a phage replication protein (A1S_1749, ACX60_RS09135) and one in a putative protein (A1S_2250, ACX60_RS06305). We hypothesized that A1S_1669 encoded a regulator of DNA Pol V genes in *A. baumannii*. The increased GFP signal of the *umuDC* gene reporter could be explained by the nonsense mutation in the middle of the ORF of the A1S_1669 gene, which would lead to a truncated protein at the codon encoding leucine 123 (L123*) (Figure 2A and B).

**Figure 2:**
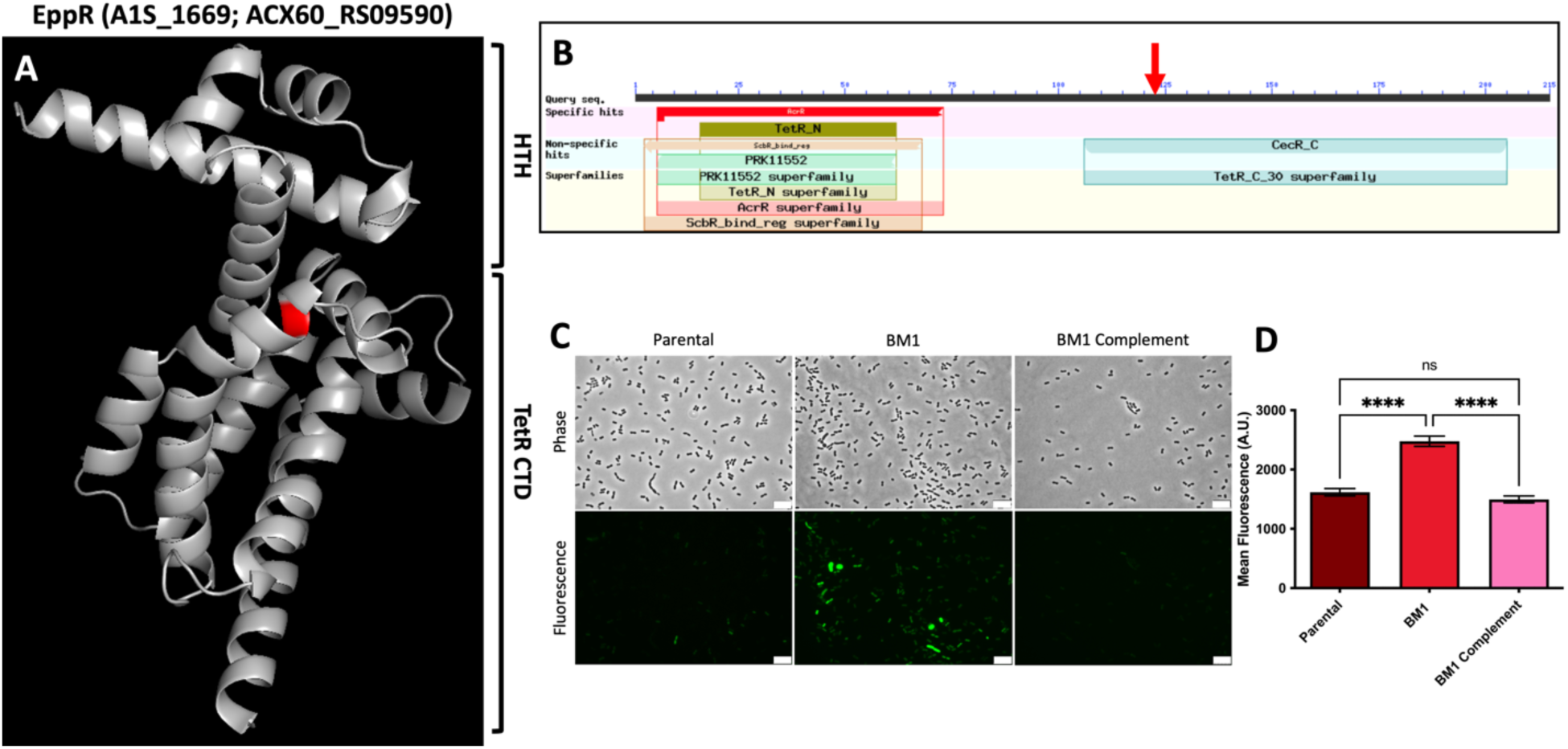
EppR (A1S_1669; ACX60_RS09590) belongs to the TetR family of bacterial transcription factors. (A) Predicted 3D-structure of EppR. The protein model of EppR was created from the primary sequence using the web-based Alphafold (Jumper et al., 2021; Mirdita et al., 2022) program and rendered using PyMOL (Schrödinger LLC, 2015) software. The model is based on known crystal structures of other TetR-like proteins. Using NCBI BLAST (Altschul et al., 2005, 1997), a helix-turn-helix (HTH) domain that is characteristic of TetR-like regulators was identified in the N-terminal domain. The C-terminal domain (CTD) is also characteristic of those found in TetR-like proteins. The location of the nonsense mutation, a mutation resulting in a stop codon, present in the *eppR* gene of the BM1 strain (L123*) is labeled in red. This model only shows a monomer, while this family of transcriptional regulators are known to dimerize. (B) Schematic of conserved domains predicted by NCBI BLAST. The nonsense mutation, shown by the red arrow, is located within the beginning of the C-terminal domain. (C) *PumuDC-gfp* shows increased expression in BM1. *umuDC*(0636-0637) expression measured by fluorescence microscopy using a *umuDC*(0636-0637) GFP transcriptional reporter (MacGuire et al., 2014) was visualized in the parental strain, BM1 mutant containing an empty pNLAC vector (BM1), and BM1 mutant complemented pNLAC containing the *eppR* gene (BM1 Complement). The white scale bar denotes 10 μm. (D) Quantification of mean fluorescence of *PumuDC-gfp*. Mean fluorescence was quantified using Fiji (Schindelin et al., 2012) and data depicted using Graphpad Prism. Compared to the parental, the BM1 mutant shows increased GFP expression. Complementation of BM1 with the *eppR* gene results in fluorescence levels like the WT and significantly lower the BM1 mutant. Approximately 1000 cells were quantified per strain. Ordinary one-way ANOVA with multiple comparisons by Tukey-test was used for statistical analysis. **** indicates a p-value of < 0.0001, n.s indicates not significant.

To confirm whether the TetR-like family regulator, which we have named EppR (<u>E</u>rror-<u>p</u>rone <u>p</u>olymerase regulator), has a role in the regulation of genes encoding DNA Pol V, we first complemented BM1 with a plasmid-borne copy of *eppR* (Table S1, Figure 2C), and measured single cell fluorescence in all the strains (Figure 2D). As expected, the fluorescence in BM1 is elevated compared to the parental strain (Figure 2C and D; BM1) while in the complemented BM1 strain (Figure 2C and D; BM1 Complement, Table S1), fluorescence is reduced to levels like the parental strain. Using Fiji (Schindelin et al., 2012), we quantified the mean fluorescence in cells of each strain. We found that in BM1, the increase in GFP signal compared to the parental strain was significant while there was no significant difference in expression comparing the parental and the complemented strains. These results suggest that the increase in *umuDC*^0636-637^ reporter expression in BM1 is due to the nonsense mutation within the *eppR* gene and not due to the other mutations present in the BM1 strain, providing evidence that EppR acts as a repressor for *umuDC*^0636-637^.

To further characterize this putative transcriptional repressor, we modeled EppR using Alphafold (Jumper et al., 2021; Mirdita et al., 2022) (Figure 2A) and searched for known conserved domains using the National Center for Biotechnology Information (NCBI) Basic Local Alignment Search Tool (BLAST) (Altschul et al., 2005, 1997) (Figure 2B). NCBI BLAST results show that EppR contains two distinct domains. The N-terminal domain (NTD) contains a helix-turn-helix (HTH) DNA binding domain that is highly conserved in TetR-like family proteins (Ramos et al., 2005; Yu et al., 2010). The C-terminal domain (CTD) is also conserved in TetR-like family proteins serving both as the ligand-sensing and dimerization domain (Ramos et al., 2005; Yu et al., 2010). The L123* mutation in the *eppR* gene present in the BM1 mutant strain is located towards the beginning of the CTD and suggests that BM1 produces a truncated variant of EppR, resulting in the derepression of *umuDC*^0636-637^.

### EppR is a Regulator of genes coding for DNA Pol V

We have shown that in BM1 a nonsense mutation in *eppR* resulted in the upregulation of one of the DNA Pol V operons, providing evidence that EppR may serve as a repressor of genes encoding DNA Pol V. *A. baumannii* contains two UmuDC operons (*umuDC^0636-0637^, umuDC^1174-^1173)* and two genes encoding UmuC (*umuC^2008^, and umuC^2015^*) (Norton et al., 2013). To test for the effect of EppR on the expression of all genes encoding DNA Pol Vs, including *umuDAb*, we created plasmid-borne GFP transcriptional reporters (Table S1) using the promoter sequences of all genes encoding DNA Pol V and *umuDAb*. We then introduced them into *A. baumannii* WT, a strain with a clean deletion of *eppR* (*ΔeppR,* Table S1), and *ΔeppR* with a plasmid-borne copy of *eppR* (*ΔeppR* Complement, Table S1). Expression was visualized using fluorescence microscopy (Figure 3A) and quantified using Fiji (Schindelin et al., 2012) (Figure 3B). We found that across all DNA Pol V genes and *umuDAb*, expression is significantly upregulated in strains lacking EppR (Fig. 3A and B). We also see that in the complementation strain, expression for all genes encoding DNA Pol V and *umuDAb* returns to levels similar or lower than WT. The lower expression of the complement is likely due to the multicopy nature of the plasmid used for complementation (Luke et al., 2010). These data are consistent with our initial experiments in the BM1 strain and show that EppR is a repressor involved in the regulation of all genes encoding DNA Pol V and *umuDAb*.

**Figure 3:**
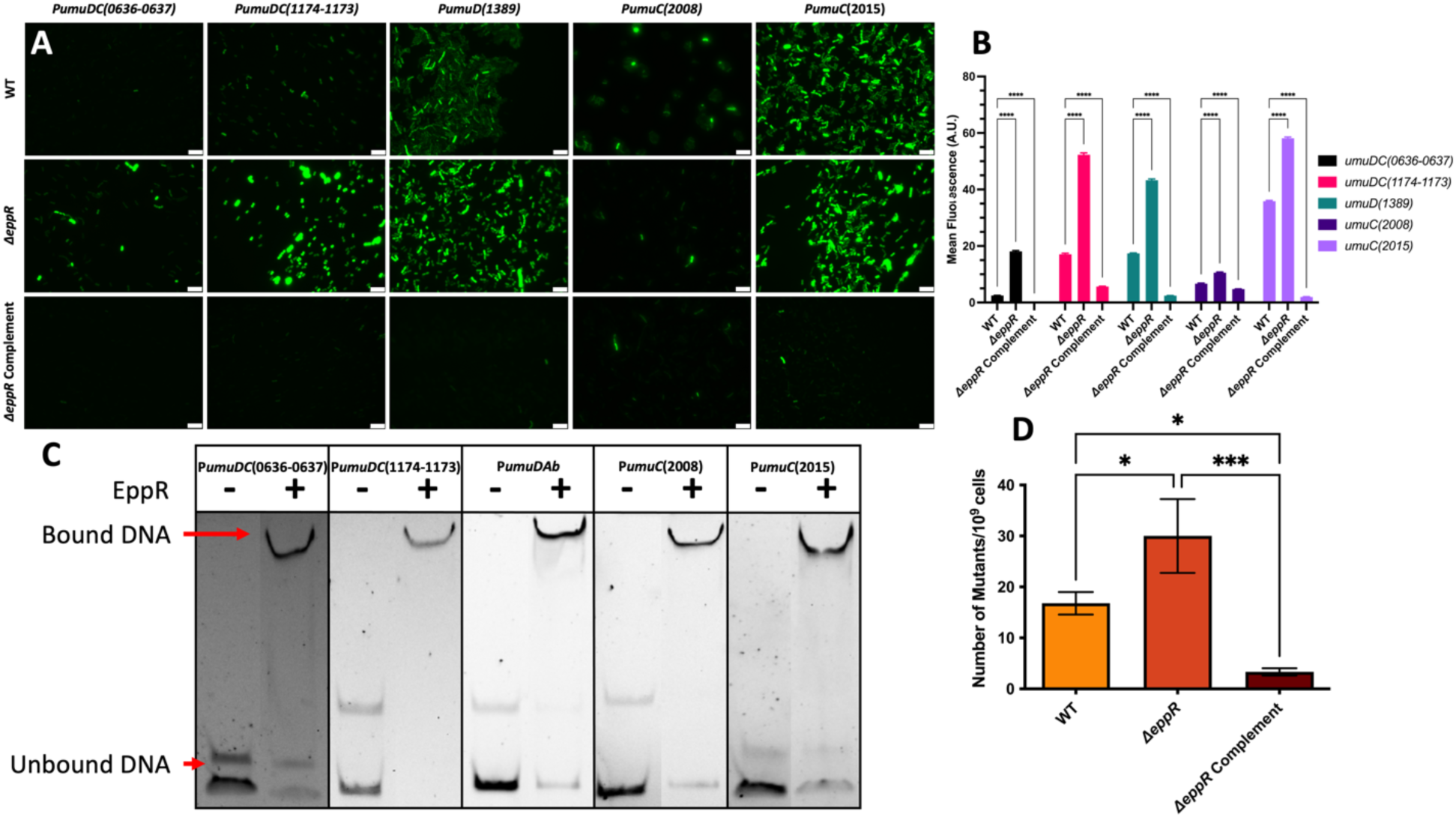
EppR regulates the expression of genes encoding DNA Pol V and *umuDAb* (A) *ΔeppR* results in increased fluorescence signal in strains with transcriptional reporters of genes encoding DNA Pol V and *umuDAb*. Expression of DNA Pol V and *umuDAb* encoding genes was visualized by fluorescence microscopy measuring the plasmid-based GFP fluorescence from the respective transcriptional reporters in WT, Δ*eppR*, and Δ*eppR* complemented with the *eppR* gene. For each strain, at least 1000 cells were visualized and quantified. The white scale bar denotes 10 μm. Corresponding phase images are provided in Figure S3. (B) Quantification of mean fluorescence of DNA Pol V and *umuDAb* reporters. The mean fluorescence measured in each of the strains with the fluorescent reporters was quantified using Fiji (Schindelin et al., 2012) and plotted with Graphpad Prism. In all cells with the fluorescent reporters, the deletion of *eppR* results in a significant increase in expression that is complemented by the *eppR* gene, indicating that EppR acts as a repressor of these genes. Ordinary one-way ANOVA with multiple comparisons by Tukey-test was used for statistical analysis. **** indicates a p-value of < 0.0001. (C) EMSA of EppR binding to a Cy3 5’-labeled DNA fragment containing the promoter region of each of the genes encoding DNA Pol V and *umuDAb*. This experiment was performed with either no EppR or 1.6 µM EppR and with 2.5 nM of Cy3-labelled DNA. Binding was observed for all promoters of genes encoding DNA Pol V and *umuDAb*. (D) *ΔeppR* results in increased frequency of spontaneous Rifampicin-resistant (RifR) mutants. EppR is a repressor of genes involved in acquisition of antibiotic resistances. To measure the effect of EppR in mutagenesis, the number of RifR mutants was determined in WT, Δ*eppR*, and the Δ*eppR* complement strains. The number of spontaneous RifR mutants were counted and standardized to 10^9^ cells. Higher frequency of RifR mutants is measured in Δ*eppR* compared to both WT and the Δ*eppR* complemented with EppR. Ordinary one-way ANOVA with multiple comparisons by Tukey-test was used for statistical analysis. * indicates a p-value of < 0.05 and *** indicates a p-value of < 0.001.

Since we have shown that EppR acts as a repressor of genes encoding DNA Pol V and *umuDAb*, we next investigated its mechanism. As NCBI BLAST (Altschul et al., 2005, 1997) (Figure 2) identified EppR as a TetR-like family transcriptional regulator, we hypothesize that EppR would act as a repressor through direct binding of DNA Pol V and *umuDAb* gene promoters. To test this, we amplified the promoter regions of the genes encoding DNA Pol V and *umuDAb* by PCR using fluorescent primers. We then performed electrophoretic mobility shift assays (EMSAs) by incubating purified EppR (Figure S1) with the fluorescence-tagged promoters and separating the products in a non-denaturing polyacrylamide gel (Figure 3C). We found that at the tested concentration of EppR (1.6 µM), EppR specifically bound to all five promoters. All EMSA experiments include an excess of unlabeled non-specific DNA competitor. The data from both the fluorescent microscopy and the EMSA experiments provide strong evidence that EppR acts as a direct repressor of genes encoding DNA Pol V and *umuDAb* in *A. baumannii*.

Since the previous experiments have shown that EppR acts as a direct repressor of genes encoding DNA Pol V and *umuDAb*, we next wanted to investigate whether EppR derepression of genes encoding DNA Pol V results in any antibiotic resistance as it has been shown that DNA Pol V activity leads to increased acquisition of antibiotic resistances (Boshoff et al., 2003; Cirz et al., 2005; Ryan T Cirz et al., 2006; Cirz and Romesberg, 2006). We performed a rifampicin mutagenesis assay in which WT, *ΔeppR*, and the *ΔeppR* complement strains were deposited on LB agar plates with or without rifampicin. Spontaneous rifampicin-resistant (RifR) mutants were counted and standardized to the number of colony-forming units (CFUs) grown on plates lacking rifampicin. The frequency of mutation was calculated as the number of RifR mutants per 10^9^ cells (Figure 3D). We observed that the mutation frequency in *ΔeppR* was approximately two times higher compared to WT. Additionally, we see that the complementing *ΔeppR* strain resulted in a mutation frequency dropping to levels below WT. These results show that EppR regulation of the DNA Pol Vs has a modest but significant effect on the acquisition of antibiotic resistances.

### Regulation of EppR and Identification of the EppR Binding Site

We have shown that EppR acts as a repressor of the genes encoding DNA Pol V and *umuDAb* through direct binding of their promoters. Since EppR is likely a TetR-like transcriptional regulator, proteins that tend to regulate their own expression (Cuthbertson and Nodwell, 2013; Deng et al., 2013; Ramos et al., 2005), we investigated if EppR binds to its own promoter by EMSA. Like the other promoters used here, we amplified the *eppR* promoter with fluorescent primers. Afterwards, we incubated the amplified promoter with varying concentrations of purified EppR (0 – 1.6 µM) and separated the reactions using a polyacrylamide gel (Figure 4A). The percent of DNA bound by EppR was quantified using Fiji (Schindelin et al., 2012).

**Figure 4:**
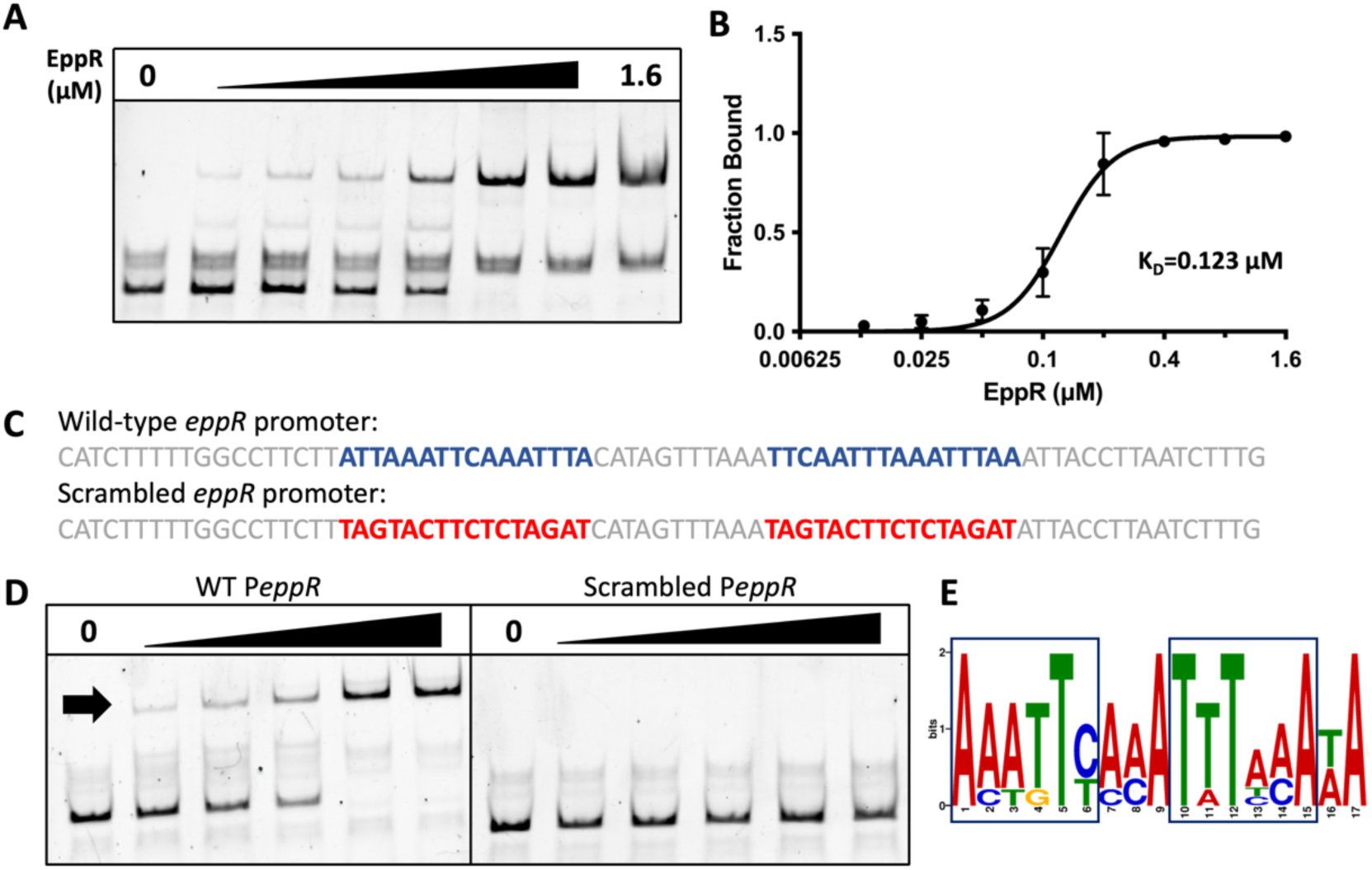
EppR binding to its own promoter permits the identification of an EppR binding motif. (A) EppR binds to a Cy3 5’-labeled DNA fragment containing the *eppR* gene promoter region. The EMSA experiment was performed with increasing EppR (0-1.6µM) and with 2.5 nM of Cy3-labelled DNA. EppR fully binds to its own promoter at concentrations as low as 0.8 µM. (B) Quantification of *PeppR* EMSA. Percentage of bound DNA was quantified using Fiji (Schindelin et al., 2012) and error bars represent the standard deviation from the analyses of three independent EMSA replicates. (C) Identification of potential EppR binding motif. A potential EppR binding site was identified (blue font) by comparing EppR binding to the site and to scrambled site sequences (red font). (D)Scrambled EppR binding motif results in loss of EppR binding. EMSAs using the native or scrambled predicted *eppR* binding site were performed to test for EppR binding at EppR concentrations ranging from 0-0.8 µM. Complete loss of EppR binding was observed in the scrambled promoter. (E) Prediction of consensus EppR binding motif. Using the Motif for Em Elicitation (MEME) program (Bailey et al., 2009), a consensus binding motif was built from the DNA Pol V, *umuDAb*, and *eppR* promoters. The conserved inverted repeat that serves as EppR binding site is demarked by the boxes.

We found that EppR binds better to its own promoter than to the DNA Pol V or *umuDAb* promoters. We begin to observe binding at concentrations as low as 0.0125µM and observed full binding at approximately 0.400 µM. The calculated dissociation constant (K_D_) of this binding is 0.123 μM (Figure 4B). This result provides evidence that EppR may regulate its own expression through direct binding of its promoter and that EppR binding is better than to the other tested promoters.

Since we have shown that EppR binds to the promoters of the genes encoding DNA Pol V, UmuDAb, and to its own promoter, we sought to determine a possible EppR binding motif, expecting this motif to be conserved in the genes that we tested. We looked first at the *eppR* promoter since EppR binding had a lower binding affinity than the *umuDC* operon promoters (Figures 3C, 4A, and 4B). It is known that TetR-family proteins tend to bind to indirect repeats (Cuthbertson and Nodwell, 2013; Ramos et al., 2005), which we identified a pair of within the *eppR* promoter (Figure 4C). To confirm that this is indeed the EppR binding site, we scrambled these sequences and performed a series of EMSA experiments to determine if there were any changes to the binding affinity (Figure 4C and D). We found that when the potential EppR binding sequence was scrambled, there was a complete loss in EppR binding. When compared to the native promoter, the upper DNA band, indicating DNA-protein binding, is absent in the EMSA (Figure 4D) with the scrambled promoter sequences at all tested concentrations of EppR. With these results, we have identified the EppR binding site within its own promoter.

Next, we wanted to assess if this sequence was also located within the promoter regions of the genes encoding DNA Pol V and *umuDAb*. Using the Motif for Em Elicitation (MEME) program (Bailey et al., 2009), we used the binding sequence from the *eppR* promoter along with the promoter sequences from the genes encoding DNA Pol V and *umuDAb* to build a consensus binding motif (Figure 4E). While we see that the *eppR* promoter contains two indirect repeats, only one of these is present in the DNA Pol V and *umuDAb* promoters.

This evidence suggests that one repeat is sufficient for EppR binding to the DNA Pol V and *umuDAb* promoters. It also suggests a mechanism to explain the tighter binding of EppR to its own promoter compared to the DNA Pol V and *umuDAb* promoters.

Next, we wanted to test whether EppR represses its own expression *in vivo*. We expected high *eppR* expression in a *ΔeppR* strain. To test this, we constructed a plasmid-borne GFP transcriptional reporter with the *eppR* promoter (Table S1) and introduced it into WT, *ΔeppR,* and *ΔeppR* complement strains. GFP fluorescence, a measurement of *eppR* expression, was visualized by fluorescence microscopy and quantified using Fiji (Schindelin et al., 2012) (Figure 5A). As expected, we found that in the *ΔeppR* strain, *eppR* expression is significantly elevated compared to the WT (Figure 5A). We also see that expression levels in the complement are reduced to levels below the WT. These results confirm that EppR represses its own expression.

**Figure 5:**
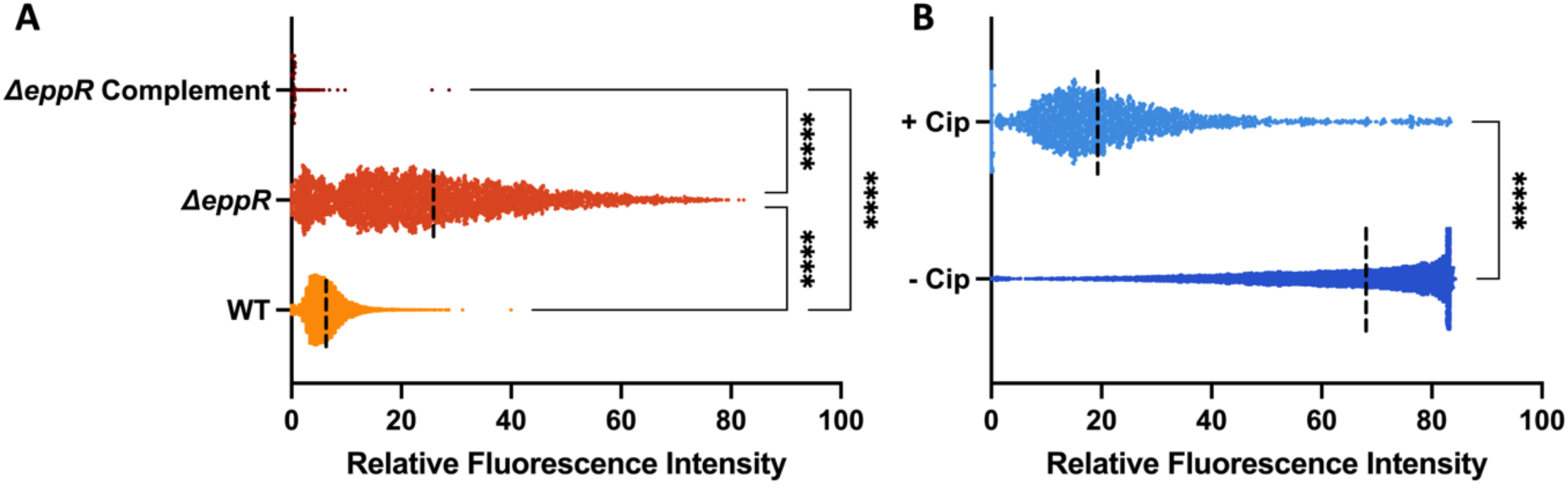
EppR regulates its own expression and is downregulated in DNA damaging conditions. (A) The deletion of *eppR* results in an upregulation of *eppR* expression. EppR regulates its own expression by acting as a repressor of *eppR* gene expression, measured in WT, Δ*eppR*, and *ΔeppR* complement strains with a P*eppR*-GFP transcriptional fluorescent GFP reporter by single cell fluorescence microscopy and quantified using Fiji (Schindelin et al., 2012). At least 1000 cells were quantified from each strain using Fiji. Ordinary one-way ANOVA with multiple comparisons by Tukey-test was used for statistical analysis. **** denotes a P-value <0.0001. (B) *eppR* expression is downregulated in response to DNA damage. WT cells treated with sublethal concentrations of Cip (+Cip, 0.2 µg/mL) show a significant decrease in fluorescence of the *eppR* GFP reporter compared to untreated cells. At least 1500 cells were quantified from each strain using Fiji. An unpaired T-test was used for statistical analysis. **** denotes a P-value <0.0001.

Next, we also wanted to investigate *eppR* expression in strains in which the DDR is induced by treatment with a DNA damaging agent. We used the antibiotic ciprofloxacin (Cip) to induce the DDR (Drlica et al., 2009; Wentzell and Maxwell, 2000). As we have shown that EppR directly represses DNA Pol V and *umuDAb* expression, we believed that *eppR* expression would be downregulated in response to DNA damage, which would then lead to derepression of genes encoding DNA Pol V and *umuDAb*. We performed fluorescence microscopy on WT cells with the *eppR* reporter treated with or without a sublethal concentration of Cip (Drlica et al., 2009; Wentzell and Maxwell, 2000), and quantified single cell fluorescent expression (Figure 5B). We found that compared to untreated cells, Cip-treated cells had approximately three-fold reduction in *eppR* expression. With these experiments, we show that EppR likely regulates its own expression through direct repression, like other TetR-family proteins, and that expression is downregulated during DNA damage. In addition, we have confirmed the EppR binding motif within the *eppR* promoter and found that it is conserved within the DNA Pol V and *umuDAb* promoters, revealing an EppR-controlled regulatory network in response to DNA damage.

### EppR coregulates DNA Polymerase V Expression with UmuDAb

Previous experiments shown here (Figure 3 and 4) provided evidence that EppR is a repressor of the DNA Pol V and *umuDAb* gene products by direct EppR binding to a conserved motif located within their promoters. Other groups have shown that one of the *umuD* genes, *umuD^1389^*, encodes a UmuD protein, called UmuDAb, with a helix-turn-helix (HTH) DNA binding domain in its amino terminal end (Aranda et al., 2014; Hare et al., 2014; Witkowski et al., 2016). It has been shown that UmuDAb is a direct repressor of its own expression, and of the other DNA Pol V encoding genes, with a characterized binding motif (Aranda et al., 2014; Hare et al., 2014; Witkowski et al., 2016). Within the promoter sequences of the DNA Pol V encoding genes, we have found that both the UmuDAb and EppR binding motifs are present, with the UmuDAb motif located closer to the translation start site while the EppR motif is further upstream. We observe that for the *umuDC^0636-0637^*, *umuDC^1174-1173^*, and *umuC^2015^* promoters, the EppR binding motif is approximately 20 bp upstream of the UmuDAb binding motif. For the *umuC^2008^* promoter, the EppR binding motif is approximately 80 bp upstream while intriguingly for the *umuDAb* promoter, there is significant overlap with the UmuDAb binding motif (Figure S2). Based on these findings, we hypothesize that EppR coregulates DNA Pol Vs gene expression with UmuDAb and since the EppR binding site is located further upstream than the UmuDAb binding site in the promoters of genes encoding DNA Pol V we believe that EppR has a secondary role in regulation to UmuDAb, which would be the primary repressor. To investigate this, we introduced each of the DNA Pol V fluorescent reporters to the WT strain and to strains lacking both *eppR* and *umuDAb* (*ΔeppRΔumuDAb*, Table S1). We compared their fluorescence to the same strain but complemented with either *eppR*, *umuDAb*, or with both *eppR* and *umuDAb*. Quantification was performed by single cell fluorescence microscopy (Figure 6). Since we hypothesize that EppR acts as a corepressor, we expected to see partial complementation, measured as elevated GFP signal in the *ΔeppRΔumuDAb* strain complemented with EppR compared to WT, though lower than the parental *ΔeppRΔumuDAb* in which there is no repression. We found that when *ΔeppRΔumuDAb* is complemented with just EppR (Figure 6; *ΔeppRΔumuDAb* + EppR), expression of all genes encoding DNA Pol V and *umuDAb* is reduced, though expression is still higher than that of WT. This indicates that EppR, by itself, cannot fully repress DNA Pol V gene expression in the absence of UmuDAb. Looking at the UmuDAb single complement strain (Figure 6; *ΔeppRΔumuDAb* + UmuDAb), DNA Pol V encoding gene expression is further reduced compared to the EppR single complement and expression levels more closely match that of WT. These results show that EppR cannot fully repress DNA Pol V expression in *ΔeppRΔumuDAb* and UmuDAb is required for full repression. This evidence supports our hypothesis that EppR and UmuDAb act as corepressors of genes encoding DNA Pol V and that UmuDAb is likely the primary repressor while EppR plays a secondary role. With these results, we propose a model for DNA Pol V regulation in *A. baumannii* where EppR and UmuDAb serve as corepressors (Figure 7).

**Figure 6:**
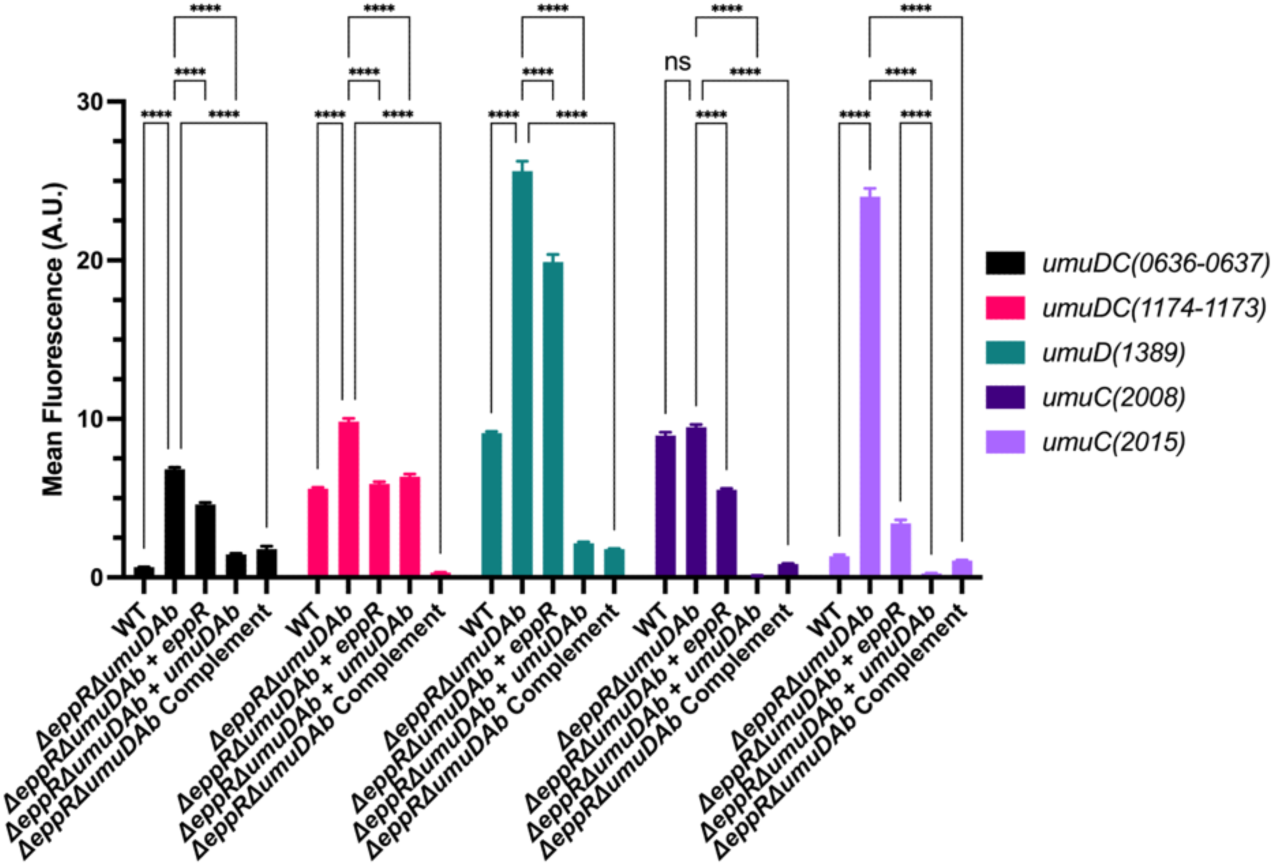
EppR and UmuDAb co-regulate DNA Pol V and *umuDAb* expression. DNA Pol V expression was measured by fluorescence microscopy using plasmid-based DNA Pol V transcriptional reporters in WT compared to the Δ*eppR*Δ*umuDAb* strain complemented with either EppR or UmuDAb, and complemented with both EppR and UmuDAb. Expression of each of the genes encoding DNA Pol V and of *umuDAb* is measured as fluorescence emitted by the respective gene reporters. EppR partially complements Δ*eppR*Δ*umuDAb* (third bar), while the full complementation is observed in the strains complemented with both EppR and UmuDAb. At least 1200 cells were quantified from each strain using Fiji (Schindelin et al., 2012). Mean GFP expression was graphed using Graphpad Prism. **** denotes a P-value <0.0001 and ns denotes no significance.

**Figure 7:**
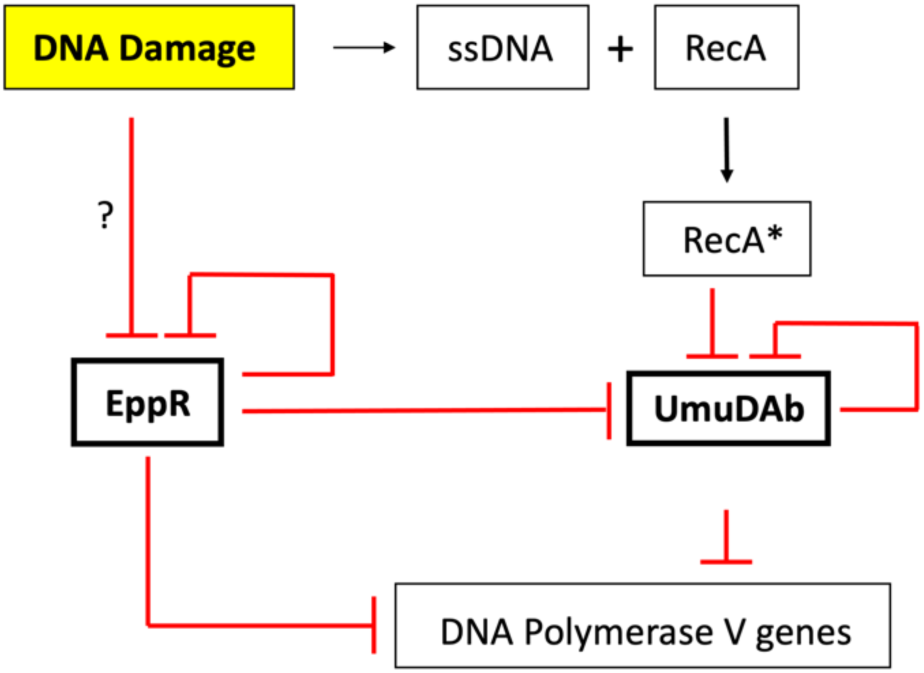
A negative circuitry involving two transcription factors regulates the expression of the genes encoding DNA polymerase V and *umuDAb* genes in *A. baumannii*. The proposed model for DNA polymerase V gene regulation includes two co-repressors, EppR and UmuDAb. In this model, both EppR and UmuDAb directly bind to the promoters of genes encoding DNA Pol V and *umuDAb* to repress their expression. In addition, EppR regulates itself through direct repression, providing a negative feedback circuitry for EppR regulation. The link between DNA damage and UmuDAb regulation of expression has been previously characterized, where the the RecA nucleoprotein filament (RecA*) interacts with UmuDAb to promote its autocleavage (Aranda et al., 2013; Hare et al., 2014, 2012; Witkowski et al., 2016). Currently, we hypothesize that EppR interacts with a ligand produced during the DDR to promote its derepression and that of the genes coding for DNA Pol Vs, this molecule remains unknown.

## Discussion

Previous work in our lab showed that there is a casual relationship between the acquisition of antibiotic resistance and the DDR (Norton et al., 2013). It has also been shown by other groups, in other bacterial species, that the activity of error-prone polymerases, namely DNA Pol V, leads to increased antibiotic resistances (Boshoff et al., 2003; Cirz et al., 2005; Ryan T Cirz et al., 2006; Cirz and Romesberg, 2006). Due to this, it is important to understand how the genes encoding these error-prone DNA polymerases are regulated in *A. baumannii*, as its DDR is not fully characterized. In this work, we provide additional insights into the regulation of genes encoding DNA Pol V and *umuDAb* in *A. baumannii*. Through a genome-wide mutagenesis assay, we have identified a novel TetR-like regulator involved with the regulation of these genes, which we have named EppR. We have found that strains lacking EppR have elevated expression levels of all genes encoding DNA Pol V as well as *umuDAb*, resulting in an approximately two-fold significantly higher mutagenesis than the parental strain (Figure 3D).

The Δ*eppR* associated phenotypes are complemented by a plasmid-borne copy of *eppR*, confirming that EppR acts as a repressor of DNA Pol V genes (Figure 3). Through EMSA, we have shown that EppR represses gene expression through the direct binding of a conserved binding motif that is present in the promoters of the genes encoding DNA Pol V and *umuDAb* (Figure 3 and 4). We also find that, like other TetR-family regulators, EppR represses its own expression through direct binding of its promoter (Cuthbertson and Nodwell, 2013; Deng et al., 2013; Ramos et al., 2005) (Figure 4 and 5). Additionally, we find that EppR expression is downregulated during the DNA damage response (Figure 5), providing a potential mechanism for the derepression of DNA Pol V genes during DNA damage, which agrees with the mutagenesis data we show here (Figure 3D). Lastly, we have shown that EppR acts as a corepressor with UmuDAb, a DNA Pol V regulator identified by other groups (Aranda et al., 2013; Hare et al., 2014, 2006a; Witkowski et al., 2016), to regulate these genes and we have proposed a model for the regulation of genes encoding DNA Pol V and *umuDAb* in *A. baumannii* (Figure 7).

Through our study, we have identified EppR as a corepressor of genes encoding DNA Pol V along with UmuDAb, which other groups have previously characterized (Aranda et al., 2013; Hare et al., 2014, 2006b; Witkowski et al., 2016). We propose a model where both proteins work together to fully repress these genes and that both will derepress during the DDR, leading to the induction of genes encoding DNA Pol V. UmuDAb expression is already well characterized and its mechanism is like that of LexA (Kelley, 2006; Little et al., 1980), where the RecA nucleoprotein filament (RecA*) interacts with UmuDAb to induce its autocleavage, leading to derepression of the genes encoding DNA Pol V (Hare et al., 2012; Witkowski et al., 2016). While we have found that EppR directly binds to the promoters of the genes encoding DNA Pol V and *umuDAb*, and that these genes are upregulated during the DDR, the mechanism of EppR-mediated derepression is still unclear. Same is true regarding the link between the induction of the DDR and EppR expression. Based on NCBI BLAST EppR is a TetR-family regulator (Figure 2). One of the characteristics of this family is that the sensor domain can bind a specific ligand, leading to a conformational change in protein structure, resulting in expression (Cuthbertson and Nodwell, 2013; Deng et al., 2013; Ramos et al., 2005). We believe that EppR-mediated expression would likely involve an interaction of the EppR sensor domain with a ligand accumulated during the DDR, though this ligand is still unknown. To gain some insights into potential ligands for EppR, we performed a homology search of the EppR sensor domain on NCBI BLAST. While it seems that EppR is unique to *A. baumannii*, we found that the EppR sensor domain shares some homology to the transcription factor RutR, which is a master regulator for the synthesis and degradation of pyrimidines (Shimada et al., 2008, 2007) It has been shown that uracil and, to a lesser extent, thymine binds to RutR as a ligand, leading to expression of the genes it represses (Nguyen Le Minh et al., 2015; Shimada et al., 2007). It is possible that EppR could utilize nucleotide bases as a ligand as well.

DDR genes are not expressed at the same time and level in response to DNA damage. The initial set of genes induced during DNA damage are those involved with high-fidelity repair, such as base excision repair (BER) and nucleotide excision repair (NER; Maslowska et al., 2019; Mullins et al., 2019; Simmons et al., 2008; Wozniak and Simmons, 2022). If these activities are unable to repair all lesions on the DNA and the DDR persists, genes encoding lower fidelity translesion DNA polymerases, including those encoding DNA Pol V, are then induced (Maslowska et al., 2019; Simmons et al., 2008). We hypothesize that during DNA damage in *A. baumannii*, BER and NER repair pathways would release nucleotide bases and if there is a large amount of damage, these bases would accumulate. As these increase in concentration, they would bind to EppR, leading to the derepression of DNA Pol V genes, which are among the last genes to be induced in a canonical DDR (Maslowska et al., 2019; Simmons et al., 2008).

When looking at the effects of co-repression of the genes encoding DNA Pol V by EppR and UmuDAb, we show that while EppR does influence the expression of these genes, UmuDAb has a greater effect on regulation. As seen in Figure 6, complementation with just UmuDAb has a stronger repressive effect compared to just EppR. This could be explained by the distance of EppR and UmuDAb binding motifs relative to the transcription start site (TSS) of the genes. It has been shown that position of the binding site of a repressor in relation to the TSS can influence how strong gene expression is repressed and can influence the mechanism of repression (Elledge and Davis, 1989; Payankaulam et al., 2010; Rojo, 1999). To gain insights into how the position of the EppR and UmuDAb binding motifs affect their ability to repress genes encoding DNA Pol V expression, we mapped the binding sites of EppR and UmuDAb in relation to the transcriptional start site (TSS) defined in Kroger et al.(Kröger et al., 2018). We found that for most of the promoters of genes encoding DNA Pol V, the UmuDAb binding motif overlaps with the TSS, where transcription initiation would likely occur, while the EppR binding site is located ∼20-80 bp upstream of the UmuDAb motif (Figure S2). The location of the UmuDAb binding site on top of the TSS is likely the reason why we and others (Aranda et al., 2013; Hare et al., 2014; Witkowski et al., 2016) believed that UmuDAb was the sole regulator of genes encoding DNA Pol V. This evidence supports our idea that UmuDAb acts as the primary regulator while EppR plays a secondary role in DNA Pol V regulation.

Our model of two proteins, and perhaps a third one (Candra et al., 2024; Cook et al., 2022, 2021; Peterson et al., 2020), regulating DNA Pol V expression contrasts with the canonical bacterial DDR, which is based oS *E. coli*, where its *umuDC* operon is repressed by a single regulator, LexA (Little and Mount, 1982; Simmons et al., 2008; Sutton et al., 2000). Having EppR as a secondary regulator may add an additional level of control to DNA Pol V expression. It is known in *E. coli* that DNA Pol V expression is tightly controlled, due to its mutagenic potential (Sutton et al., 2000). This is especially important in *A. baumannii*, since unlike *E. coli* where there is a single *umuDC* operon, there are multiple copies of *umuDC* in the genome (two *umuDC* operons and two *umuC* copies) (Norton et al., 2013). These additional copies of genes encoding DNA Pol V may necessitate an additional repressor to ensure tight regulation. This corepression model could also provide a more sophisticated level of regulation of the genes encoding DNA Pol V as expressions levels could be modulated depending on which protein is binding to the promoter. If both UmuDAb and EppR are present, we would expect fully repression of these genes. If only UmuDAb is present, we would have low levels of expression, while we would see higher levels if only EppR is present as we hypothesized that EppR plays a secondary role to UmuDAb. Lastly, if both proteins are derepressed, we would then have full expression of the genes encoding DNA Pol V. The ability to finely modulate the expression of these genes could allow *A. baumannii* to control mutagenesis levels. Intriguingly, mutagenesis is significantly though slightly elevated in the Δ*eppR* strain while *umuDAb* expression is induced (Figure 3D). The data suggest that *A. baumannii* controls the expression of genes encoding DNA Pol V in a nuanced manner; some level of mutagenesis may offer a selective advantage.

Despite the differences in the regulation of DNA Pol V in *A. baumannii* compared to the canonical DDR, the timing of gene expression during the DDR is somewhat similar. In a canonical DDR, the timing of gene expression is provided for the LexA binding motif (Courcelle et al., 2001; Fernández de Henestrosa et al., 2000; Maslowska et al., 2019; Simmons et al., 2008). The promoters of early expressing DDR genes have a weaker LexA binding motif compared to late expressing genes in the regulon. This simple but effective mechanism allows for the expression of only the early DDR genes during lower levels of DNA damage while the later genes with higher mutagenic potential, including genes encoding DNA Pol V, are only expressed if DNA damage persists (Maslowska et al., 2019; Simmons et al., 2008). We do see that EppR has the potential to act in a similar manner, due to the difference in strength of EppR binding to its own promoter versus the promoters of the genes encoding DNA Pol V. We observed that EppR binds to its promoter with higher affinity (Figures 3 and 4). This is likely both due to the presence of an additional EppR binding motif in the *eppR* promoter (Figure S2) and differences in the motif sequence in the promoters of the genes encoding DNA Pol V. During DNA damage, this would allow DNA Pol Vs to be expressed before EppR. Once *eppR* is expressed and DNA damage is resolved, the amount of EppR protein would exceed the amount of ligand needed for expression, leading to EppR rebinding the *eppR* promoter and the promoters of genes coding for DNA Pol Vs and UmuDAb, fully repressing their expression once again.

Here, we have identified a novel regulator involved in the regulation of genes encoding DNA Pol V and *umuDAb* in *A. baumannii* and have shown that the regulation involves both EppR and UmuDAb to ensure tight repression and perhaps allow a low level of expression of the error prone DNA Pols. Interestingly, it is likely that EppR may also regulate genes that are outside of the DDR. One characteristic of TetR-family regulators is that their promoters can be located between two divergent genes and control their expression, with one of the genes being the regulator itself (Cuthbertson and Nodwell, 2013). The *eppR* promoter is organized in the same way, where on one side of the promoter is the *eppR* gene and on the other (A1S_1670-1673; ACX60_RS09585, ACX60_RS09580, ACX60_RS09575, ACX60_RS09570) there is an operon putatively encoding secretion and membrane proteins, currently uncharacterized. Based on this organization, it is likely that EppR controls the expression of these genes, which do not seem to be part of the traditional DDR regulon. It would be interesting to investigate if EppR is involved in the regulation of cellular processes outside of the DDR.

## Experimental Procedures

### Strains and culture conditions

All bacterial strains (Table S1) were grown in LB medium at 37°C, with shaking at 225 rpm. Antibiotics were added to the medium at the indicated concentrations unless it is stated otherwise. Kanamycin (Km, 35 µg/mL), carbenicillin (Carb, 100 µg/mL), erythromycin (Ery, 50 µg/mL) and tetracycline (Tet, 6 µg/mL).

### Strain and Plasmid Construction

Construction of the P*umuDC^0636-0637^* GFP transcriptional reporter strain was previously described in Macguire et al.(2014).

To construct the *ΔeppR* strain, 1000 bp of the flanking DNA regions of *eppR* (A1S_1669, ACX60_RS09590) were amplified with the primer pairs EppRupF/EppRupR and EppRdownF/EppRdownR. Additionally, a kanamycin resistance cassette from a previous plasmid, pUAC66(MacGuire et al., 2014), was amplified with the primers KanFw and KanRev. The three fragments were then ligated into pUC19 (New England Biolabs). Afterwards, the fragment containing the flanking regions, and the antibiotic marker was amplified with EppRupF and EppRdownR and introduced by transformation into *A. baumannii* ATCC 17978 via electroporation, as previously described(MacGuire et al., 2014). Recombinants were selected for by kanamycin resistance and verified by PCR using the EppRupF and EppRdownR primers.

The *ΔeppRΔumuDAb* unmarked strain was constructed by amplifying and cloning 1000 bp of the flanking regions of *umuDAb* (A1S_1389) with the primer pairs 1389upF/1389upR and 1389downF/1389downR in pUC19. The desired fragment lacking the *umuDAb* gene ORF and promoter was amplified by PCR and cloned in pLGB36 (Ito et al., 2020). The plasmid was then introduced by transformation in the *ΔeppR* strain selecting for EryR colonies. This strain contains a mutated and native copy of the gene of interest. The double recombinant containing only the mutated copy and the loss of backbone from the chromosome was induced with anhydrous tetracycline (aTC)-induced toxin counterselection (ss-Bfe3). Colonies are EryS. The deletion was corroborated by PCR using the 1389upF and 1389downR primers.

To complement the BM1 mutant strain we constructed the pNLAC *PeppR-eppR*, pBN3, encoding the *eppR* gene and its promoter (Table S1). This was constructed with a fragment consisting of the *eppR* gene, and its native reporter that was amplified from genomic DNA using the primers EppRSalIF and EppRNotIR (Table S2). PCR fragments and the pNLAC vector(Luke et al., 2010) were digested with SalI and NotI (New England Biolabs) and ligated with T4 DNA ligase (Invitrogen). The mix was introduced by transformation into chemically competent *E. coli* DH5α and selected for TetR. Plasmids were sequenced using the primers PNLACF (Table S2) to verify correct assembly. The plasmid was then introduced by transformation into *A. baumannii*.

pNLAC *PumuDAb-umuDAb* (Table S1; pBN11) and pNLAC PeppR-eppR *PumuDAb-umuDAb* (pBN12) were constructed by amplifying a fragment consisting of the *umuDAb* gene and its native promoter from genomic DNA using the primers umuD(1389)_Full_BsaI_F and umuD(1389)_Full_BsaI_R (Table S2) and digested with BsaI. PCR fragments were cloned into BsaI sites present in pNLAC or pBN3 (Table S1), respectively, using Golden Gate cloning as previously described (Brychcy et al., 2023). Plasmids were introduced by transformation in *E. coli* DH5α and selected for by TetR. Plasmids were sequenced using primer PNLACF to verify correct assembly. The plasmid was then introduced by transformation into *A. baumannii*.

DNA Pol V plasmid-based transcriptional reporters (Table S1; pBN5-10) were constructed using Golden Gate cloning as described previously(Brychcy et al., 2023). Promoters of genes encoding DNA Pol V were amplified with their respective primers (Table S2) and cloned in pMB1-A along with pET ribosomal binding site (RBS), enhanced green fluorescent protein (eGFP), and B0012 terminator(Brychcy et al., 2023).

To purify EppR in *E. coli* BL21 AI with pET11t *eppR*, the *eppR* gene was amplified from genomic DNA using the primers EppRBamHI and EppRNotI (Table S2). Appropriate flanking restriction enzyme sites and a 6x histidine tag was added to the 3’ end of the gene. PCR fragments were digested with BamHI and NotI (New England Biolabs) and ligated with T4 DNA ligase (Invitrogen). Ligated products were introduced by transformation into *E. coli* DH5α and selected for by ampicillin resistance, the marker present in pET11T(Nguyen et al., 1993). Constructs were confirmed by sequencing using the PET11TF and PET11TR primers (Table S2). The plasmid was then introduced into chemically competent *E. coli* BL21 AI. The EppR protein was purified as shown below.

### Methyl methanesulfonate treatment and screen of bright fluorescent mutants

*baumannii* P*umuDC^0636-0637^* GFP transcriptional reporter strain was grown to saturation and diluted 1:50 into 10 mL of LB broth containing kanamycin. Cells were grown for 1.5 hours at 37°C, after which they were treated with 25 mM methyl methanesulfonate (MMS) for 1 hour. Cells were washed twice with PBS to remove MMS. These cells were deposited on plates of LB Agar with kanamycin and incubated overnight at 37°C. About 10,000 colonies were screened using a fluorescent stereoscopic microscope. 18 brightly fluorescent colonies with no DNA damaging agent were selected as bright mutants (BM). To ensure that changes to fluorescence intensity were not due to mutations in the reporter construct or *umuDAb*, both were amplified by PCR from the genome and sent to Eurofins for Sanger sequencing.

### Whole genome sequencing

Genomic DNA was isolated using the DNeasy Blood and Tissue Kit (Qiagen). Samples were then blunted and 3’ A tailed using the Quick Blunting Kit and Klenow Fragment (New England Biolabs). TruSeq indexed and universal adapters (Integrated DNA Technologies) were ligated to sample fragments using the Quick Ligation Kit (New England Biolabs). DNA fragments were separated on a 2% agarose gel and fragments from 600-800 base pairs were isolated using the QIAquick Gel Purification Kit (Qiagen). Fragments were then amplified by 15 cycles of PCR and sequenced using Illumina sequencing at the Tufts Core Facility (Boston, MA).

### Fluorescence Microscopy

Strains were grown in LB broth until mid-exponential phase was reached from saturated cultures. 2 µL of cell culture was placed on a 1% agarose in water pad and covered with a coverslip before imaging. Cells were visualized using a Leica MicroStation 5000 with a Leica DM3000G camera (Leica). Mean fluorescence was quantified using Fiji(Schindelin et al., 2012).

### EppR protein structure prediction and conserved domain analysis

Structure of EppR was predicted using Alphafold(Jumper et al., 2021). The model was rendered using PyMOL software. Conserved domains were found using the NCBI conserved domain database (NCBI).

### Purification of EppR

*E. coli* BL21 AI with pET11t *eppR* was used to purify EppR. Cells were grown at 37°C until saturation. This culture was used to inoculate 1.5 L of LB medium supplemented with ampicillin and 0.05% (v/v) glucose and grown at 37°C until mid-exponential phase (∼3 hr.). Protein overexpression was induced by adding 0.05% (v/v) L-arabinose. The culture was then incubated at 20°C for approximately 12 hr. Cells were spun down and lysed using a cell homogenizer at ∼14,000 psi (Stansted SPCH-10). EppR was purified by immobilized metal affinity chromatography (IMAC) using an AKTA fast protein liquid chromatography (FPLC) system (General Electric) using conditions described before(Cafarelli et al., 2013). Eluted protein was analyzed using SDS-PAGE to determine purity.

### Electrophoretic Mobility Shift Assay (EMSA)

DNA probes were created by PCR amplification with primers labelled with a Cy3 fluorophore on the 5’ end (Table S2). DNA (2.5 nM) and protein were mixed with binding buffer (10mM Tris-HCl [pH 8], 10mM HEPES, 50mM KCl, 1mM EDTA, 100 ng of salmon sperm DNA, 0.1 mg of bovine serum albumin per mL). Binding reactions were incubated for one hour at 37°C in a thermocycler (Eppendorf). Reaction mixtures (20 uL) were loaded into a 5% non-denaturing Tris-borate-EDTA (TBE) polyacrylamide gel. Binding complexes were separated at 150V for one hour at 4°C. Cy3-labelled DNA-protein complexes were visualized using a Bio-Rad Chemidoc imager (Bio-Rad). All EMSA experiments were repeated three times and quantified using Fiji(Schindelin et al., 2012).

### Identification of the EppR binding motif

Promoter regions of *eppR* and *umuDC* genes upregulated in the *ΔeppR* strain were analyzed for binding motifs using the Multiple Em for Motif Elicitation (MEME) program(Bailey et al., 2009). Potential binding sites were confirmed by scrambling the site and performing EMSA.

## Abbreviations

DDR: <u>D</u>NA <u>d</u>amage response
EMSA: <u>E</u>lectrophoretic <u>m</u>obility <u>s</u>hift <u>a</u>ssay
EppR: <u>E</u>rror-<u>p</u>rone <u>p</u>olymerase regulator
MMS: <u>M</u>ethyl <u>m</u>ethane<u>s</u>ulfonate
TSS: <u>T</u>ranscription <u>s</u>tart <u>s</u>ite
WT: <u>W</u>ild-type; *Acinetobacter baumannii* ATCC17978

## Acknowledgements

V.G-C was funded by a stipend from Northeastern University Skills for Capacity and Inclusion, an Inclusive Excellence award from HHMI The EcoFlex kit was a gift from Paul Freemont (Addgene Kit #1000000080) We would like to thank the Chai and Geisinger Lab at Northeastern University for their support, feedback, and equipment. We would like to thank the Malamy Lab at Tufts Medical School in Boston for providing us with the pLGB36 plasmid.

## Author Contributions

**Brian Nguyen:** Conceptualization, Formal Analysis, Investigation, Methodology, Validation, Visualization, Writing – Original Draft Preparation

**Carly Ching:** Formal Analysis, Investigation, Methodology

**Ashley Macguire:** Formal Analysis, Investigation, Methodology

**Pranav Casula:** Investigation, Methodology

**Connor Newman:** Investigation, Methodology

**Faith Finley:** Investigation, Methodology

**Veronica G Godoy:** Conceptualization, Funding Acquisition, Project Administration, Supervision, Writing – Original Draft Preparation

## Abbreviated Summary

A novel TetR-like regulator (EppR) has been identified to repress genes encoding DNA polymerase V in *Acinetobacter baumannii* through the direct binding of a conserved EppR motif in their promoters. EppR works with previously identified regulator UmuDAb to serve as corepressors of genes encoding DNA polymerase V. During the DNA damage response, both regulators would unbind from their promoter, though different mechanisms, resulting in the production of DNA Polymerase V.

