## Supplemental Information for "Multiple transcription factors regulate the expression of genes for error prone DNA polymerases in *Acinetobacter baumannii*"

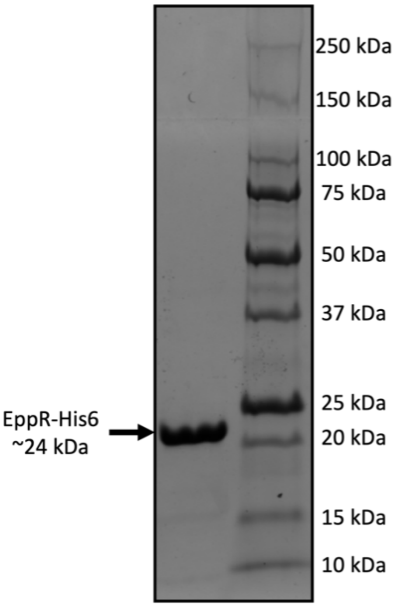


**Figure S1: SDS-PAGE of purified EppR-His6**

EppR-His6 was separated on a 12% polyacrylamide gel after purification. Approximately 30 µg of purified protein was loaded into the gel. EppR-His6 migrates between the 20 and 25 kDa size marker which is consistent with the predicted size of EppR-His6 (~24 kDa). The ladder used was Precision Plus Protein™ Unstained Protein Standards from Bio-rad.

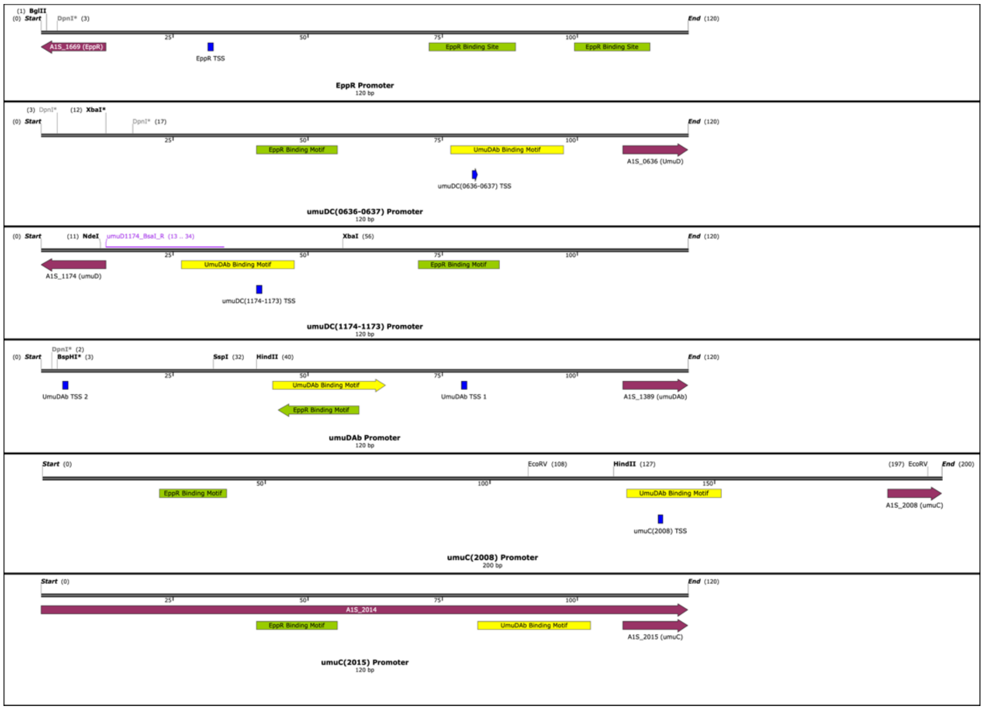


**Figure S2: Promoter Maps of genes encoding DNA Pol V, *umuDAb*, and *eppR***

The promoter maps show the organization of the EppR binding motif (green), the UmuDAb binding motif (yellow), and the predicted transcription start site (TSS)(blue)(Kröger et al., 2018).

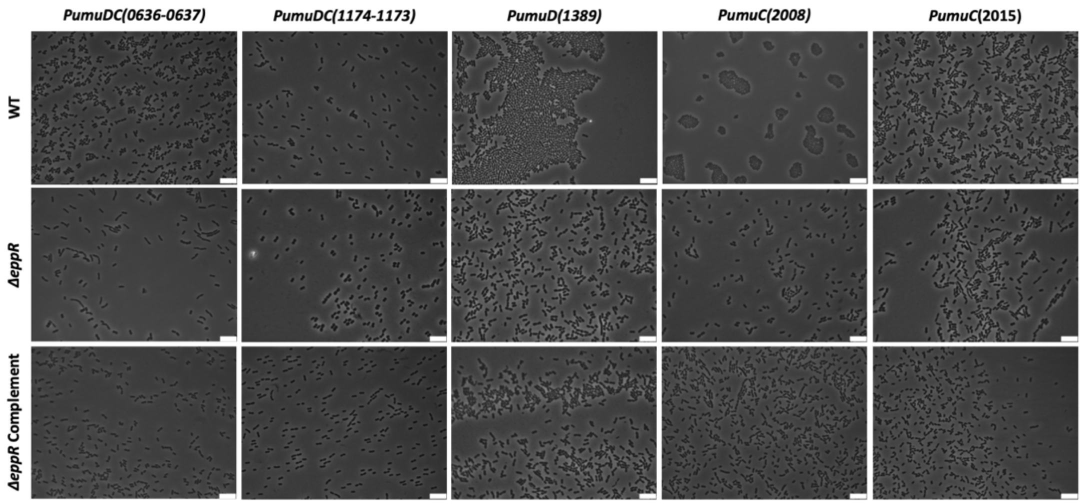


**Figure S3: Phase images for Figure 3A.**

| Strain Name or Plasmid | Genotype or Description | Reference |
|---|---|---|
| ***Acinetobacter baumannii*** | | |
| *A. baumannii* WT | ATCC17978 wild-type strain | America Type Culture Collection |
| *A. baumannii umuDC*-GFP | ATCC17978 w/ chromosomal *umuDC(0636-0637)* GFP transcriptional reporter (KanR) | Macguire et al., 2014 |
| BM1 (Bright Mutant 1) | ATCC17978 *A1S_1669 L123** (TTA→TAA)*, A1S_1748 R135G* (CGT→GGT)*, A1S_2250 F64I* (TTT→ATT) | This work |
| *A. baumannii ΔeppR* | ATCC17978 *eppR::kan* (KanR) | This work |
| *A. baumannii ΔeppRΔumuDAb* | ATCC17978 *eppR::kan* w/ unmarked *umuD(1389)* deletion (KanR) | This work |
| | ***Escherichia coli*** | |
| *E. coli* DH5α | F- *endA1 glnV44 thi-1 recA1 relA1 gyrA96 deoR nupG Φ80dlacZΔM15 Δ(lacZYA-argF)U169, hsdR17(rK- mK+)*, λ– | |
| *E. coli* BL21 AI | $F^-$*ompT hsd*$S_B$ ($r_B^-$, $m_B^-$) *gal dcm ara*B::T7RNAP-*tet*A | |
| **Plasmids** | | |
| pET11T | *E. coli* expression plasmid | |
| pNLAC | *A. baumannii* shuttle vector used for complementation (TetR, AmpR) | Luke et al., 2010 |
| pUC19 | Plasmid used to construct *eppR* and *umuDAb* knockout construct | |
| pLGB36 | Suicide vector for allelic replacement (AmpR, EryR) | Ito et al., 2020 |
| pMB1-A | Golden Gate Plasmid w/ BsaI sites that replicates in *A. baumannii* (AmpR) | Brychcy et al., 2023 |
| pBN1 | pET11T *eppR*-His6 (AmpR) EppR purification | This work |
| pBN2 | pLGB36 w/ *umuD(1389)* flanking regions (AmpR, EryR), UmuDAb knockout plasmid | This work |
| pBN3 | pNLAC *PeppR-eppR* (TetR), EppR complementation plasmid | This work |
| pBN4 | pNLAC w/ AmpR counterselection gene removed (TetR) | This work |
| pBN5 | pMB1-A w/ *PumuDC(0636-0637)* (ACX60_18290; ACX60_18295)-GFP transcriptional reporter (AmpR) | This work |
| pBN6 | pMB1-A w/ *PumuDC(1174-1173)* (ACX60_RS12360; ACX60_RS12365)-GFP transcriptional reporter (AmpR) | This work |
| pBN7 | pMB1-A w/ *PumuDAb*-GFP transcriptional reporter (AmpR) | This work |
| pBN8 | pMB1-A w/ *PumuC(2008)* (ACX60_RS07740)-GFP transcriptional reporter (AmpR) | This work |
| pBN9 | pMB1-A w/ *PumuC(2015*(ACX60_RS07700)*)*-GFP transcriptional reporter (AmpR) | This work |
| pBN10 | pMB1-A w/ *PeppR* (ACX60_RS09590)-GFP transcriptional reporter (AmpR) | This work |
| pBN11 | pNLAC *PumuDAb-umuDAb)* (TetR), UmuDAb complementation plasmid | This work |
| pBN12 | pNLAC *PeppR-eppR PumuDAb)-umuDAb* (TetR), EppR and UmuDAb complementation plasmid | This work |

### Table S1: Strain and Plasmid List

| Oligonucleotides | Sequence 5' to 3' | Description |
|---|---|---|
| EppRupF | CTTGGGCAGACGCAAGTTGA | Used to construct *ΔeppR* strain |
| EppRupR | AGCTGGCAATTCCGACGTCTATCACTTCGTCTAGATCTGG | Used to construct *ΔeppR* strain |
| EppRdownF | TCGCTTGGACTCCTGTTGATTGCTCAGATGCCATTACATG | Used to construct *ΔeppR* strain |
| EppRdownR | GAACAATTAGCACCGCAGTTA | Used to construct *ΔeppR* strain |
| KanFw | CTATGGTACCAGACGTCGGAATTGCCAGCT | Used to construct *ΔeppR* strain |
| KanRev | CTATGGTACCTTACTGTCCCTAGTGCTTGG | Used to construct *ΔeppR* strain |
| EppRHis6_F | GGGGGCATATGATGTCCAGATCTAGACGAAGTG | Used to construct pBN1 |
| EppRHis6_R | TTTTTGGATCCCTAATGATGATGATGATGATGTCCTCCTCCTCCTCCATT TTTAAGATTTGATTGGCTAATTGCTTTTAATCC | Used to construct pBN1 |

| | | |
|---|---|---|
| pET11TF | CCCCAAGGGGTTATGCT | Used to verify pBN1 |
| pET11TR | TACGACTCACTATAGGGGAATTGT | Used to verify pBN2 |
| umuD(1389)up_SalI_F | CTAAGTCGACAGCGTTGAGTGTGTATGAATAGG | Used to construct pBN2 |
| umuD(1389)up_KpnI_R | CTATGGTACCATCGCCTCCATTTCACCGTTC | Used to construct pBN2 |
| umuD(1389)down_KpnI_F | CTATGGTACCCCTATAACCTCAAACGAATGAGAT | Used to construct pBN2 |
| umuD(1389)down_BamHI_R | GCAAGGGATCCCGTTCCTTGTATCAGCAGG | Used to construct pBN2 |
| EppRSalIF | GCGCATCGTCGACCTTGGGCAGACGCAAGTTGA | Used to construct pBN3 |
| EppRNotIR | GTTAGCGGCCGCGAACAATTAGCACCGCAGTTA | Used to construct pBN3 |
| PNLACF | AGTTTGCGCAACGTTGTTGCCA | Used to verify pNLAC-based constructs |
| PNLACR | AACGACGAGCGTGACACCAC | Used to verify pNLAC-based constructs |
| PumuD0636_BsaI_F | ATTCATATGGGTCTCACTTAGCTACAACCACGTTTTAATTTT | Used to construct pBN5 |
| PumuD0636_BsaI_R | ATTGCATGCGGTCTCAGTACATTAAAACCTCTTGAAAACGTAACG | Used to construct pBN5 |
| PumuD1174_BsaI_F | ATTCATATGGGTCTCACTATGACAAAAGGCAAGACTATACTTAC | Used to construct pBN6 |
| PumuD1174_BsaI_R | ATTGCATGCGGTCTCAGTACATGTTTCCCCTAGCTTGATTTT | Used to construct pBN6 |
| PumuD1389_BsaI_F | ATTCATATGGGTCTCACTATTTCGCATGTCTCCATACCA | Used to construct pBN7 |
| PumuD1389_BsaI_R | ATTGCATGCGGTCTCAGTACATCGCCTCCATTTCACC | Used to construct pBN7 |
| PumuC2008_BsaI_F | ATTCATATGGGTCTCACTATAAAAAACTTAAGCGTATTATAAAAAAAGCTGT | Used to construct pBN8 |
| PumuC2008_BsaI_R | ATTGCATGCGGTCTCAGTACTATTTCATCCCCTTACTCATTAACTG | Used to construct pBN8 |
| PumuC2015_BsaI_F | ATTGCATGCGGTCTCAGTACTATTTCATCCCCTTACTCATTAACTG | Used to construct pBN9 |
| PumuC2015_BsaI_R | ATTGCATGCGGTCTCAGTACACCATTTTGAAATCGTAACAAATTC | Used to construct pBN9 |
| PeppR_BsaI_F | ATTCATATGGGTCTCACTATCTTTTTGGCCTTCTTATTAAATTCAAATTTAC | Used to construct pBN10 |
| PeppR_BsaI_R | ATTGCATGCGGTCTCAGTACTGTTTAAACCTAATTTTATTTCATCATAAAAGTATATCT | Used to construct pBN10 |
| umuD(1389)_Full_BsaI_F | ATTCATATGGGTCTCACTATTTCGCATGTCTCCATACCA | Used to construct pBN11 and pBN12 |
| umuD(1389)_Full_BsaI_R | TATGGTCTCATCTGGTCAGTAACTTAAATAATAAAATATAAGAAAAT | Used to construct pBN11 and pBN12 |
| MB21 | ATTTCAGATAAAAAAAATCCTTAGCTTTCG | Used to verify pBN5-10 |
| MB31 | TGCAGCGTCTCTGGCA | Used to verify pBN5-10 |
| P0636_EMSA_F | TGAATGAAGCTATAAGTAAAATTG | Used to amplify EMSA fragment |
| P0636_EMSA_R_Cy3 | AATAAAGTAGAGCACAGCTT | Used to amplify EMSA fragment |
| P1174_EMSA_F | ATTTGCTATAAATAAGCAATAGTGTGGAACATTAAG | Used to amplify EMSA fragment |
| P1174_EMSA_R_Cy3 | ATGTTTCCCCTAGCTTGATTTTGTAACAT | Used to amplify EMSA fragment |
| P1389_EMSA_F | ACTCATATTTAAGTGTTCAGAATTTCATTG | Used to amplify EMSA fragment |
| P1389_EMSA_R_Cy3 | ATCGCCTCCATTTCACC | Used to amplify EMSA fragment |
| P2008_EMSA_F | GCATTACTACATTTAAAAGACTCAACTCA | Used to amplify EMSA fragment |
| P2008_EMSA_R_Cy3 | TATTTCATCCCCTTACTCATTAACTGAAA | Used to amplify EMSA fragment |
| P2015_EMSA_F | CTAGATCAGAACTGAAGAATTTCCGAC | Used to amplify EMSA fragment |
| P2015_EMSA_R_Cy3 | ACCATTTTGAAATCGTAACAAATTCAAG | Used to amplify EMSA fragment |
| PeppR_EMSA_F_Cy3 | CTTTTTTATTCATCTTTTTGGCCTTCTTATTAAATTC | Used to amplify EMSA fragment |
| PeppR_EMSA_R | GGACATGTTTAAACCTAATTTTATTTCATCATAAAAG | Used to amplify EMSA fragment |

**Table S2: Oligonucleotide List**
